# Bacterial community profiles and *Vibrio parahaemolyticus* abundance in individual oysters and their association with estuarine ecology

**DOI:** 10.1101/156851

**Authors:** Ashley L. Marcinkiewicz, Brian M. Schuster, Stephen H. Jones, Vaughn S. Cooper, Cheryl A. Whistler

## Abstract

Oysters naturally harbor the human gastric pathogen *Vibrio parahaemolyticus*, but the nature of this association is unknown. Because microbial interactions could influence the accumulation of *V. parahaemolyticus* in oysters, we investigated the composition of the microbiome in water and oysters at two ecologically unique sites in the Great Bay Estuary, New Hampshire using 16s rRNA profiling. We then evaluated correlations between bacteria inhabiting the oyster with *V. parahaemolyticus* abundance quantified using a most probable number (MPN) analysis. Even though oysters filter-feed, their microbiomes were not a direct snapshot of the bacterial community in overlaying water, suggesting they selectively accumulate some bacterial phyla. The microbiome of individual oysters harvested more centrally in the bay were relatively more similar to each other and had fewer unique phylotypes, but overall more taxonomic and metabolic diversity, than the microbiomes from tributary-harvested oysters that were individually more variable with lower taxonomic and metabolic diversity. Oysters harvested from the same location varied in *V. parahaemolyticus* abundance, with the highest abundance oysters collected from one location. This study, which to our knowledge is the first of its kind to evaluate associations of *V. parahaemolyticus* abundance with members of individual oyster microbiomes, implies that sufficient sampling and depth of sequencing may reveal microbiome members that could impact *V. parahaemolyticus* abundance.

## 1 Introduction

Shellfish, including the eastern oyster (*Crassostrea virginica*), are common vectors for human pathogens. This includes the bacterium *Vibrio parahaemolyticus*, the leading causative agent of bacterial seafood-borne gastroenteritis worldwide and an emergent pathogen in the United States (US) (1). Oysters concentrate *V. parahaemolyticus* from overlaying water, which can lead to a naturally higher abundance than the < 10,000 Most Probable Number (MPN)/g that is currently recommend in the US as a limit to ensure shellfish is safe for consumption (2, 3). To increase shellfish safety, fisheries employ strategies intended to reduce pathogen levels in live product, including depuration in UV sterilized water or relay/transplantation of oysters to a location where *V. parahaemolyticus* is of low abundance, often correlating with high salinity (4, 5, 6). But reported correlations of salinity with *V. parahaemolyticus* abundance from environmental studies are mixed (3) suggesting factors other than salinity likely mediate a reduction in *V. parahaemolyticus* concentrations. Furthermore, relay of oysters into non-sterilized water more effectively reduces *V. parahaemolyticus* contamination than depuration in sterile water (6). Therefore, antagonistic relationships among community members coupled with less than favorable salinity conditions could explain the greater reduction of *V. parahaemolyticus* in the oyster microbiome during relay (7, 8).

Ecological studies reveal a few biotic factors correlate with *V. parahaemolyticus* abundance both in water and in oysters (3). Zooplankton can positively correlate with *V. parahaemolyticus* abundance when they serve either as a nutrient resource or a mechanism of dispersal (3, 9). *V. parahaemolyticus* abundance also positively correlates with chlorophyll *a*, suggesting a general interaction with phytoplankton (3, 8). Oysters could passively accumulate planktonic and particle-associated Vibrios by filter-feeding (3). Even so, the overall oyster microbiome is more diverse than the overlying water microbiome, suggesting potential selective accumulation and culturing of some microorganisms, including Vibrios, by the oyster (10-14). *in vitro* bacterial-*Vibrio* competitions illustrate several types of marine bacteria influence *Vibrio* abundance, suggesting *in situ* interactions could influence accumulation in oysters (15-17). Although a growing number of studies have profiled the oyster and overlying water microbiome (18-23), none have yet attempted to correlate presence or abundance of species or community composition profiles to the relative abundance of *V. parahaemolyticus*.

To identify the core and variable microbiome among individual oysters, and correlate differences in *Vibrio parahaemolyticus* abundance with microbiome composition, we profiled the microbiome of individual oysters and overlying water from two naturally occurring, ecologically-distinct oyster beds. We employed 454 pyrosequencing of 16s RNA variable region (V2-V3) amplicons, and in parallel quantified *V. parahaemolyticus* abundance. We determined that oysters harbor a microbiome that is distinct from overlying water, and species composition and relative abundance is influenced by location of the oyster bed, potentially reflective of unique ecology. The abundance of *V. parahaemolyticus* varied between oysters and correlated with only a few rare phyla, which were linked to location. The study suggested increased sampling and microbial community sequencing depth could reveal meaningful patterns of association both with ecology and potentially *V. parahaemolyticus* abundance.

## 2. Results and Discussion

### 2.1 Sequencing the oyster microbiome

To assess the composition of and variation in oyster associated microbiota, native oysters were collected from two ecologically distinct sites, less than five miles apart in the Great Bay Estuary (GBE) of New Hampshire. The Oyster River (OR) oyster bed is located within one of the seven tributaries of this estuary where harvesting is prohibited due to its proximity to the outflow of a municipal wastewater treatment facility (WWTF), whereas the Nannie Island (NI) oyster bed is centrally located within the estuary and is classified as approved for recreational harvesting (8). Thus, these two sites may reveal how the different associated ecological and sewage discharge-related factors pertaining to each site influence the microbial community composition. We generated and sequenced 16s rRNA gene amplicons from total bacterial DNA isolated from ten individual oyster homogenates and one overlying water sample from each collection site. From the generated ∼ 1.5 million reads, only ∼ 1/3 (512,220) with 100% identity to the forward primer and mid-tag were included in the analysis. Quality filtering with FlowClus removed an additional 6,995 reads, producing an average of 18,087 reads per NI oyster (10,338-31,788; n=10), and 29,391 reads per OR oysters (ranging from 9,670-47,231; n=10) for analysis. A lower number of reads were obtained from water – 6495 and 397 from NI and OR, respectively (Supplemental Table 1). Because we seek to determine correlations of identifiable phylotypes with estuarine conditions and *V. parahaemolyticus* abundance, we evaluated in parallel two common pipelines, QIIME and mothur, that use different clustering algorithms, and determined mothur maximized assignment of OTUs at the species level of classification and also resulted in no unclassified reads (Table 1). Therefore, analysis continued with the mothur-generated classified dataset.

**Table 1.**
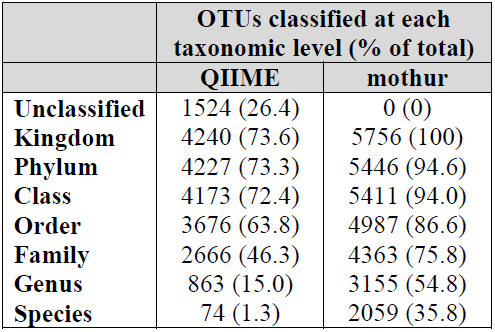
OTU assignment to level of taxonomic classification following processing.

The rarified phylogenetic distance (PD) whole tree alpha diversity index was applied to illustrate within-sample diversity and evaluate sufficiency in depth of sequencing. Overall higher index values in NI samples indicated higher alpha diversity than OR samples (Fig. 1). However, he plotted rarefactions demonstrated that total phylogenetic distances between all OTUs at each subsampling step continued to increase with higher sampling, indicating that all interpretations should consider that the data did not capture total diversity (Fig. 1).

**Figure 1.**
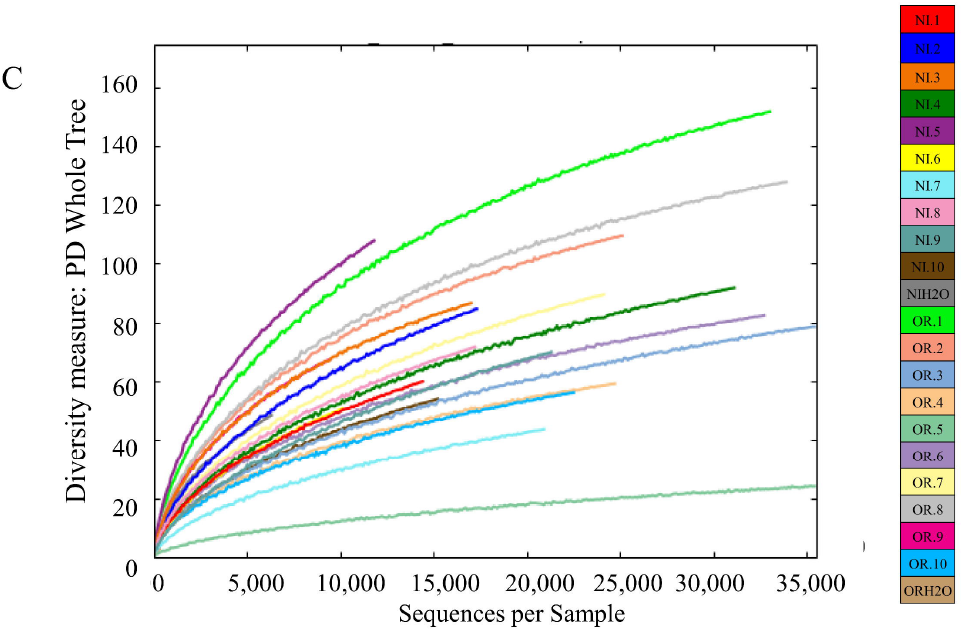
Rarified alpha diversity assessment. Phylogenetic Distance (PD) whole tree index for individual oyster and overlying water samples were generated where each sample is represented by a unique color.

### 2.2 Comparison of the distribution of phylotypes by site and substrate

Comparisons of the distribution of OTUs in individual oysters and overlying water can reveal the extent to which the microbiome of individual oysters reflects the microbes in overlying water. Since relatively fewer reads were available from the water samples, it is unsurprising that oysters from both sites harbor OTUs absent in the respective overlying water, (Fig. 2A). Even so, some OTUs from the water samples (1.35% and 0.19% from NI and OR, respectively) were not detected in any oyster, suggesting the potential for some selectivity in the microbiome accumulated from water, which is in agreement with other oyster microbiome studies (18, 25).

**Figure 2.**
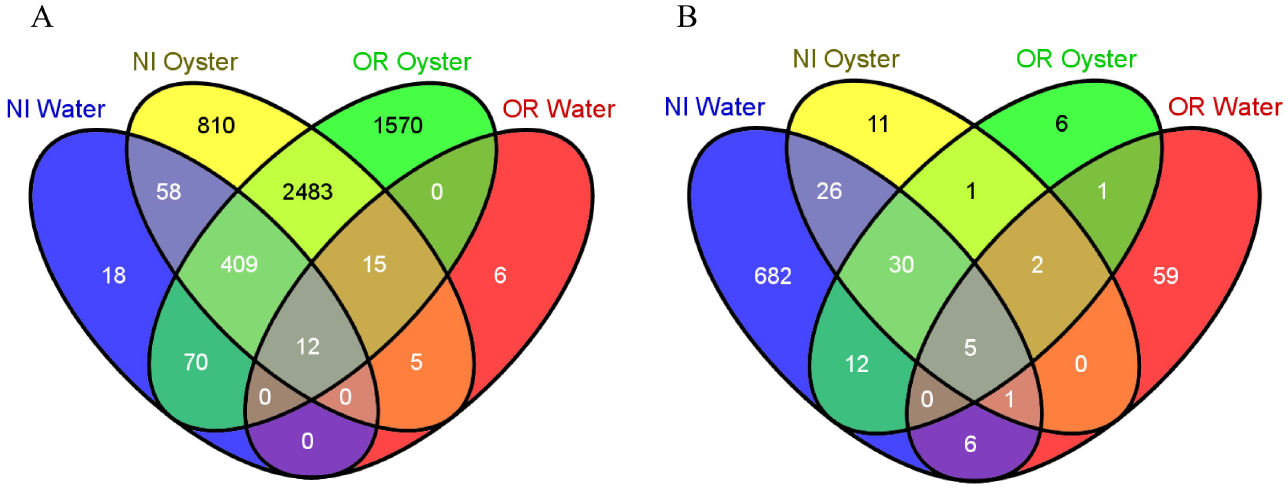
Relationships between OTUs identified in oysters and overlying water from two estuarine locations. (A) Distribution of all OTUs in water and in any oyster sample and (B) distribution of all OTUs in water and in every oyster sample by site (OR or NI), representing the site-specific and overall core microbiome.

Comparisons of individual oysters to each other also provided insight into the shared and variable microbiome. Less than 1% of the OTUs identified in any oyster were present in every one of the 20 oysters. However, the shared microbiome from oysters harvested from the same location was slightly larger, with 2.29% OTUs shared between every oyster from NI, and 1.25% shared from OR (Fig. 2B). In addition, 82% of the OTUs shared between every NI oyster were also present in overlying water (1.63% of total OTUs present in any NI oyster), indicating these consistently detected microbiome members at this site are substantially present in and likely influenced by the water column. Because the OR water sample yielded so few sequences for analysis, meaningful comparisons in this case were deemed not possible.

Next, we evaluated whether there were informative patterns in the abundance and distribution of phyla-level classifications by dual hierarchical cluster analysis, representing broad scale differences between the microbiome of both sampling sites. Most of the variation between sample type, sites, and even individual oysters was explained by not the high abundance, but the mid-and low abundance phyla (Fig. 3A), and rarifying the sequences to remove the lowest-abundant OTUs, which is a common practice to remove erroneous OTUs (e.g., 20), would have removed most of this potentially informative variation. Phylum-level analysis clearly demonstrated differences between the overlying water and oysters. This analysis also revealed delineation between the microbial communities of oysters by site, when considering both standardized and unstandardized clustering, with only a few exceptions (Fig. 3A-B).

**Figure 3.**
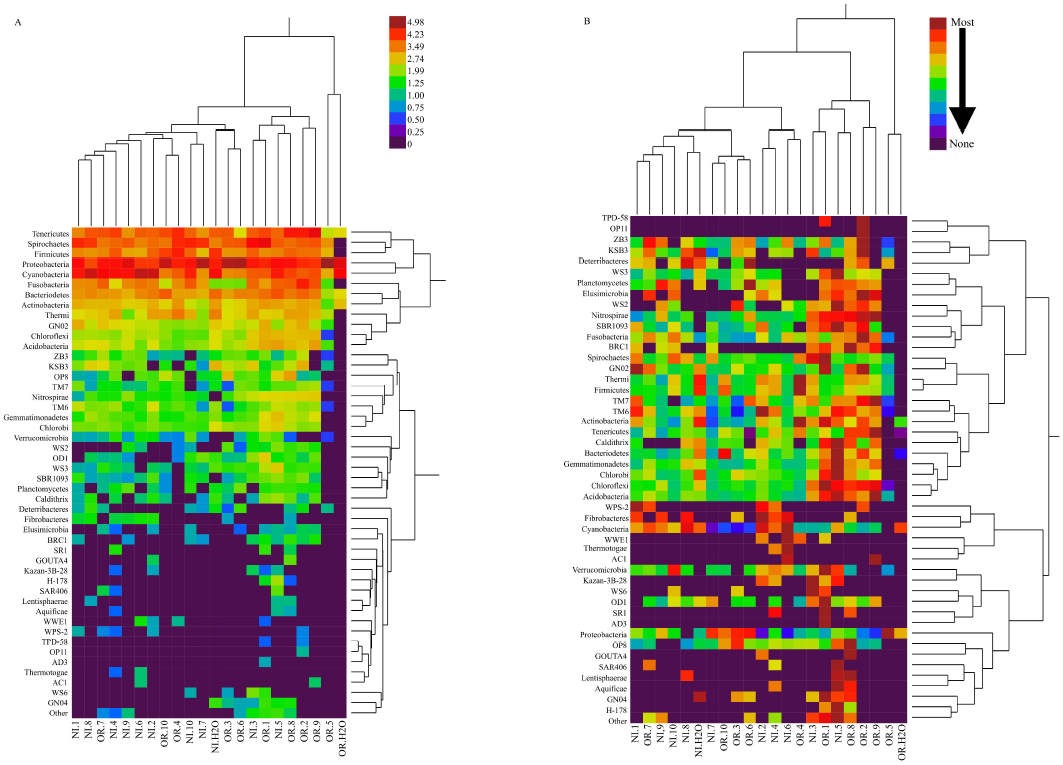
Dual hierarchical analysis of phyla-level classification. The log-transformed percent abundance of each phylum is indicated by a color scale. Samples and phyla are clustered based on (A) unstandardized and (B) standardized average linkage. In unstandardized linkage, the abundance of each phylum in a given sample is colored based on relative abundance of all phyla, whereas in standardized the abundance of each phylum is colored based on the relative abundance of that phylum across all samples.

The most abundant phyla were consistent with other oyster microbiome studies, even though these were from relatively warmer climates. The major *Crassostrea* sp.-associated phyla include Cyanobacteria, Chloroflexi, Firmicutes, Proteobacteria, Planctomyces, and Bacteriodetes (18, 20, 21). The digestive gland of Sydney rock oysters contains many of these same phyla, and also is dominated by Spirochaetes (23). Cyanobacteria were in higher abundance at NI (33.8%, ranging from 1 to 69%) than OR (7.7%, ranging from 0.8 to 43.0%; Fig. 3B). Whereas some oyster microbiome studies have discarded Cyanobacteria reads to eliminate sequenced chloroplasts from algal matter (20, 21), oysters will ingest Cyanobacteria as a food source (26) and accumulate Cyanobacteria in greater numbers than the surrounding water column (18), justifying retention of these reads as part of the microbiome. Cyanobacteria may even influence the abundance of other members of the oyster microbiome. For instance, Proteobacteria, Bacteriodetes, and Firmicutes have all been isolated from cyanobacterial blooms (27). Therefore, it is possible that differences in microbial community composition between NI and OR were influenced, at least in part, by the overall higher abundance of Cyanobacteria at NI.

Whereas differential abundances in broad phyla-level classifications reveal general patterns, considering all taxonomic levels with Unifrac uncovered more specific relationships between samples (Fig. 4). Unifrac delineated between sampling sites, with only a few exceptions. NI oysters clustered together, whereas OR oysters were dispersed among several branches, indicating NI oyster microbiomes, which had overall fewer unique phylotypes than OR, are overall more similar to each other than are OR oyster microbiomes. Unifrac analysis revealed variance (see NI.7) that was not apparent in the dual-hierarchical clustering which resulted from a classification level deeper than phylum. The OR water sample, which had the lowest read coverage, was quite distant from most samples in both clustering analyses, with a proportionally high number of reads assigned to the genus *Octadecabacter* (43.3%, compared to the average of 0.2% for all other samples, ranging from 0.05% to 0.5%) and the Mamiellaceae family (40.8%, compared to the average of 0.1% for all other samples, ranging from 0.002 to 0.5%).

**Figure 4.**
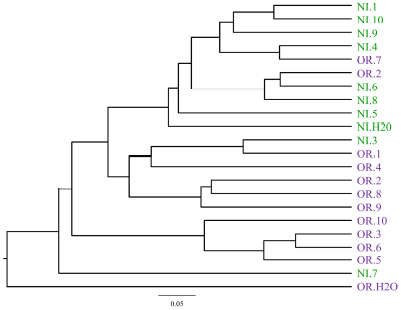
Unifrac phylogenetic distance analysis. Phylogenetic distance of all taxonomic levels for each pair of samples was calculated by the total branch length of unique phylotypes and divided by total branch length of all phylotypes.

The apparent differences in the oyster microbiome between the two sampling sites were further interrogated by employing LEfSe to identify phylotypes that significantly differ by site. The proportions of four phylotypes were significantly higher in OR oysters than NI oysters including *Finegoldia*, *Bradyrhizobium*, *Roseateles depolymerans*, and *Brevundimonas intermedia* (Fig. 5). In contrast, the proportions of eight phylotypes were significantly higher in NI oysters compared to OR including *Propionigenium*, M2PT2_76, *Reinekea*, *Pseudomonas viridiflava*, *Clostridium sticklandii*, *Vibrio fortis*, *Halobacillus yeomjeoni*, and Endozoicimonaceae. *Finegoldia* is typical of the human gastrointestinal tract (28) and *Bradyrhizobium* is a soil-dwelling, root nodule organism (29), so these associations with OR are consistent with site being a narrow tidal tributary where the oyster bed is in close proximity to the terrestrial environment and a WWTF outfall. Conclusions on associations by site of the other organisms are not possible due to absence of relevant information in published studies.

**Figure 5.**
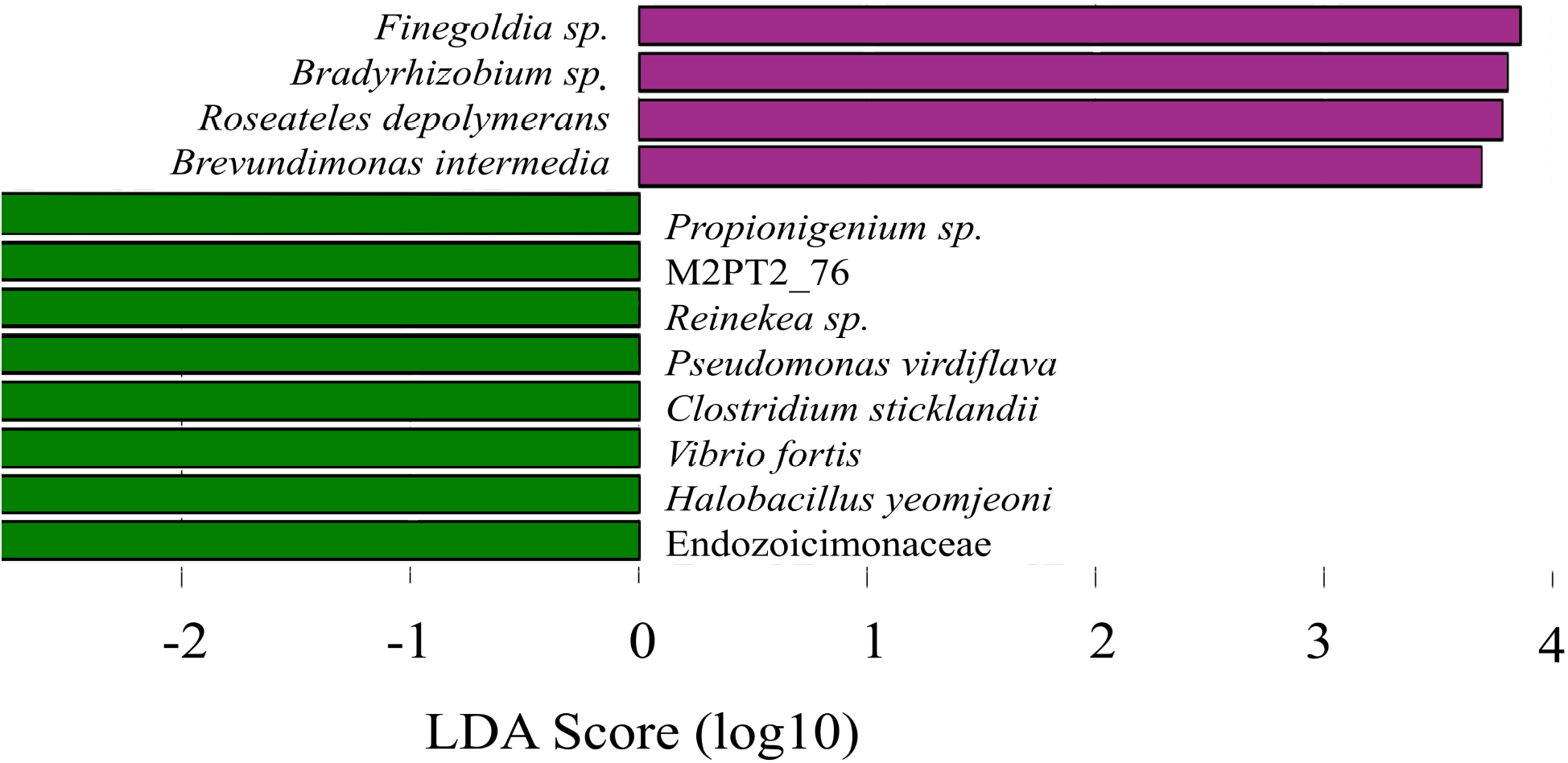
Effect size of phylotypes in oysters at significantly different proportions at each collection site determined by LEfSe. LEfSe employs the non-parametric factorial Kruskal-Wallis sum-rank, Wilcoxon rank-sum, and Linear Discriminant Analysis tests to determine the effect size of significantly different phylotypes (56).

### 2.3 Differences in predicted functional profiles between sites

In addition to defining the members of the oyster microbiome, we investigated potential functional differences inferred from phylotype composition between the sampling sites, which may be driven by their unique ecological and environmental associations. For this we used the bioinformatics tool PICRUSt that draws upon previously sequenced genomes and annotations (30). A total of 887 predicted gene functions significantly differed between NI and OR oyster microbiomes (p < 0.05). Further examination of the functional differences of the 11 functions at p < 0.0005 reveals two distinct classes (Fig. 6). OR oyster microbiomes had a higher number of functions generally involved in cell growth, including nucleotide metabolism, tRNA synthesis and associated elongation factors, amino acid biosynthesis, and oxidative phosphorylation. NI oyster microbiomes had a higher number of diverse metabolic functions (sugar, chlorophyll, carbon, and sulfur metabolism) as well as higher number of chaperone-associated proteins. The more diverse photosynthesis related metabolic capacity logically related to the prevalence of 16s sequences identified as Cyanobacteria at NI, as compared to OR. Overall, the variations between microbiomes and their respective predicted functions at each site could relate to nutrient conditions that support different types of organisms. The OR oyster bed not only is impacted by a nearby municipal WWTF discharge, which could explain the slightly higher levels of dissolved nutrients associated with WWTF effluent (orthophosphate, nitrate), but also more directly influenced by rainfall/runoff events and nonpoint source pollution (Table 2). OR has higher chlorophyll *a*, water temperature, and turbidity with lower salinity and dissolved oxygen compared to NI (Table 2A&B). NI is also a much larger oyster bed with abundant oyster cultch on coarser textured sediment compared to OR. The chronic loading of readily available nutrients at OR may support more rapid total growth of a less diverse bacterial population, whereas the lower, potentially limiting nutrient concentrations available at NI may support a more diverse bacterial population, in agreement with the measures of alpha diversity (Fig. 1). This pattern of overall lower taxonomic and function diversity in OR is in agreement with other studies that indicate wastewater effluent decreases microbial diversity (31, 32).

**Table 2.**
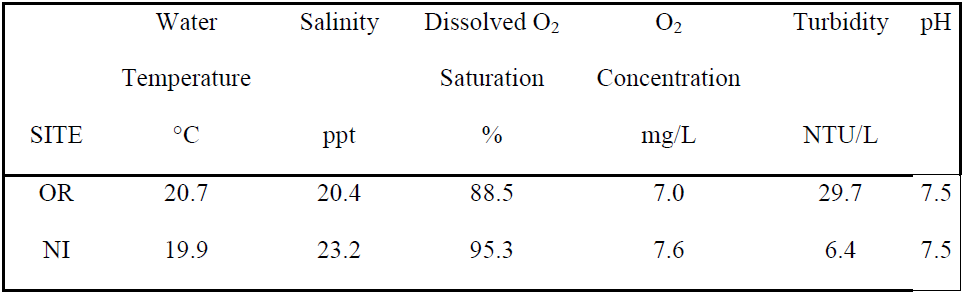
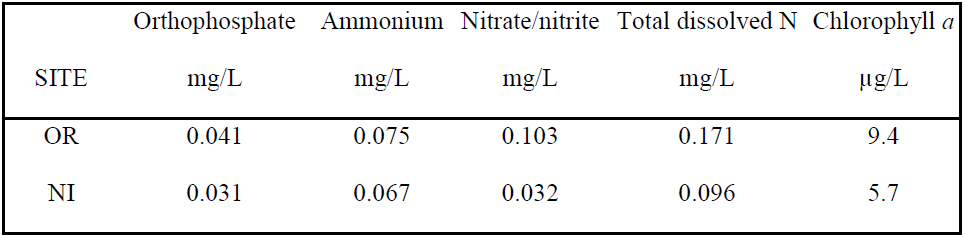
Environmental conditions associated with collection sites A. Average datasonde ^1^ measures during 12 hours prior to sampling on 8/31 to 9/1/09. B. Average concentrations in grab samples collected during July-August during 2007-2009.

|  | Water | Salinity | Dissolved O<sub>2</sub> | O<sub>2</sub> | Turbidity | pH |
| --- | --- | --- | --- | --- | --- | --- |
|  | Temperature |  | Saturation | Concentration |  |  |
| SITE | °C | ppt | % | mg/L | NTU/L |  |
| OR | 20.7 | 20.4 | 88.5 | 7.0 | 29.7 | 7.5 |
| NI | 19.9 | 23.2 | 95.3 | 7.6 | 6.4 | 7.5 |

B. Average concentrations in grab samples collected during July-August during 2007-2009.
|  | Orthophosphate | Ammonium | Nitrate/nitrite | Total dissolved N | Chlorophyll *a* |
| --- | --- | --- | --- | --- | --- |
| SITE | mg/L | mg/L | mg/L | mg/L | µg/L |
| OR | 0.041 | 0.075 | 0.103 | 0.171 | 9.4 |
| NI | 0.031 | 0.067 | 0.032 | 0.096 | 5.7 |

<sup>1</sup>Data derived from a YSI datasonde deployed at each site with readings taken at 15 min intervals. Ppt: parts per thousand; NTU: Nephelometric Turbidity Unit.

**Figure 6.**
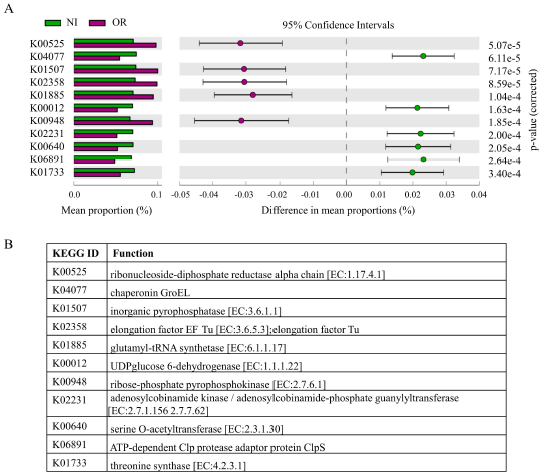
Distribution of Annotated functions at each site. (A) PICRUSt-derived KEGG orthology IDs at significant different (p < 0.0005) numbers at each site, and (B) the pathways associated with each ID.

### 2.4 Abundance of *V. parahaemolyticus* in individual oysters and correlations with microbiome

To evaluate potential correlations of microbiome with *V. parahaemolyticus*, we applied a quantitative qPCR-based MPN enumeration method of *V. parahaemolyticus* to individual oysters (see methods), which allowed evaluation of correlations between relative abundance of *V. parahaemolyticus* and phylotypes in individual oyster microbiomes. This approach revealed that individual oysters, even from the same site, differed dramatically in abundance of *V. parahaemolyticus* (Table 3) as reported by two other individual oyster studies (32, 34). Oysters were subsequently categorized and grouped based on log10 MPN/g abundance level, where the means of each group significantly differed from the other groups (Low: 0.48, Medium: 1.16, High: 2.51; p < 0.0001). *V. parahaemolyticus* was only captured via 16s sequencing in the medium and high abundance level oysters from NI (Table 3), being an overall rare component of the sequenced oyster microbiome. Whereas the estimated relative (16s) and absolute (MPN) *V. parahaemolyticus* abundance did not match, there was agreement in the general pattern of detection of *V. parahaemolyticus* 16s rRNA and abundance by enrichment-based MPN.

**Table 3.** Distribution of *Vibrio parahaemolyticus* in oysters as determined by MPN and 16s sequencing^1^.

| Oyster | Log10 MPN/g | Abundance Level | 16s Reads |
| --- | --- | --- | --- |
| NI.10 | 0.52 | Low | - |
| NI.5 | 0.72 | Low | - |
| NI.7 | 1.01 | Medium | - |
| NI.8 | 1.01 | Medium | 2 (0.011) |
| NI.1 | 1.34 | Medium | 3 (0.020) |
| NI.6 | 1.34 | Medium | - |
| NI.2 | 2.38 | High | - |
| NI.3 | 2.38 | High | 2 (0.011) |
| NI.4 | 2.38 | High | 1 (0.003) |
| NI.9 | 2.88 | High | 1 (0.009) |
| OR.10 | 0.13 | Low | - |
| OR.4 | 0.28 | Low | - |
| OR.6 | 0.28 | Low | - |
| OR.2 | 0.49 | Low | - |
| OR.8 | 0.52 | Low | - |
| OR.5 | 0.93 | Low | - |
| OR.1 | 1.01 | Medium | - |
| OR.3 | 1.01 | Medium | - |
| OR.7 | 1.20 | Medium | - |
| OR.9 | 1.34 | Medium | - |

<sup>1</sup>Oysters from Nannie Island (NI.1-10), and Oyster River (OR.1-10) are ordered by site and within site by increasing *V. parahaemolyticus* abundance level as determined by MPN. 16s reads represents the number of *V. parahaemolyticus* sequences and relative percent abundance in parentheses

NI harbored the only oysters with high abundance level of *V. parahaemolyticus*, and also harbored oysters with medium and low abundance level. In contrast, OR contained oysters with only medium and low abundance level *V. parahaemolyticus*. Due to these differences in distribution of *V. parahaemolyticus* abundance, we queried whether differences in ecology between these sites could impact microbial communities including Vibrios. There are some differences in long-term nutrient conditions (Table 2; 35) between the two sites. The combination of higher chlorophyll *a*, turbidity, nutrients (Table 2) and temperature, with lower salinity and dissolved oxygen levels (representing short-term environmental conditions averaged over the 12 hours prior to oyster harvest) at OR are consistent with it being a tributary and other distinctions between the two sites (Table 2A&B). Interestingly, although chlorophyll *a* positively correlates with *V. parahaemolyticus* presence even in the GBE (3, 8), it was higher at OR. It is not clearly apparent that any of these measured abiotic parameters drove higher levels of *V. parahaemolyticus* at NI in a subset of oysters, or comparatively lower levels of *V. parahaemolyticus* at OR, but it is further evidence supporting site differences as a likely contributing factor in oyster microbial community variation.

To investigate whether microbial community members correlated with *V. parahaemolyticus* abundance, microbiome data for individual oysters were analyzed with Unifrac distance trees to determine similarity of the microbiome of oysters in the same MPN abundance level group, separated by site. Branching patterns did not correspond with *V. parahaemolyticus* abundance level, indicating there is no overall similarity in the microbial community in *V. parahaemolyticus* high abundance level oysters. Despite a lack of clustering of samples by *V. parahaemolyticus* abundance level, there were 24 phylotypes significantly higher in number in high abundance level oysters, one phylotype in medium abundance level oysters, and three in low abundance level oysters (Fig. 7). However, a caveat to this data and its interpretation is that these were all rare phylotypes of which the proportion could be influenced by depth of sequencing of the microbiome (Fig. 1), in addition to the relationships potentially being confounded by site-specific differences. Specifically, the 19 phylotypes that were exclusive to high abundance level oysters (and by default present in significantly higher proportions) could be an artifact of oyster location in the estuary. As such, the influence of estuary location on differences in the microbiome was most apparent from this study, and correlations of microbiome with *V. parahaemolyticus* abundance await a more robust sample size and greater microbiome sequencing depth.

**Figure 7.**
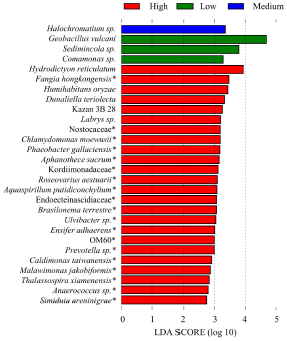
PICRUSt-derived phylotypes in oysters at different proportions by *V. parahaemolyticus* abundance class. Phylotypes followed by an ^*^ were only present in high abundance oysters.

## 3. Conclusions

Microbial community profiling of the microbiome and quantification of *Vibrio parahaemolyticus* abundance of oysters and overlaying water from two naturally occurring oyster beds revealed individual oysters and sites have different taxonomic and functional microbiome profiles. These differences, likely influenced by distinctive ecology, is in general agreement with other studies that conclude the microbiomes of marine animals are highly specific based on individuals’ surrounding habitat (23, 25) and diet (36). Even so, that the microbiome composition and Vibrio abundance were so variable between individuals from the same site allude to the potential that community-level interactions within an oyster impact the risk of *Vibrio parahaemolyticus* achieving an infective population size. A better understanding of these interactions could open new avenues for disease prevention.

Both culture-based and culture independent methods revealed *V. parahaemolyticus* did not equally accumulate in individual oysters, despite the oyster's exposure to the same general environmental conditions at each site. Therefore, the measured environmental conditions cannot explain differences in *V. parahaemolyticus* levels between individual oysters. Bivalves actively filter water based upon particle size (37), bacterial species (38), strains within the same species with known or introduced (i.e., mutations) genetic variation (39-41), and even differentiate between viral particles (42). The *V. parahaemolyticus* strains themselves may contain genetic factors or phenotypic traits influencing uptake and/or depuration and it is possible different strains are accumulated at different rates, much like *Vibrio vulnificus* (43). Through differential killing of strains or variants, oyster hemocytes may also reduce the accumulation of certain *V. parahaemolyticus* strains (44, 14).

The higher abundance of Cyanobacteria at NI may influence the abundance of other phyla at this site (27). It may also explain the higher abundance of *V. parahaemolyticus* at NI. Cyanobacteria and *V. cholerae* will associate (3), and *Vibrio* spp. make up as much as 6% of all cultivable heterotrophic bacteria isolated from cyanobacterial blooms (27). In addition, cyanobacterial-derived organic matter increases *Vibrio* abundance (3). This study indicates the general approach of microbiome profiling may reveal phylotypes and functional differences associated with *V. parahaemolyticus* abundance with deeper sampling.

## 4. Methods

### 4.1 Oyster Collection and Processing

One water and ten oyster samples were collected at low tide on September 1^st^, 2009 from two distinct naturally-occurring oyster beds in the New Hampshire Great Bay Estuary (GBE). Oysters were collected using oyster tongs whereas water samples were collected by submerging capped sterile bottles ∼ 0.5m below the water surface and uncapping to fill. Samples were immediately stored on ice packs in coolers until laboratory processing. Individual oysters were cleaned, aseptically shucked, and thoroughly homogenized with a surface disinfected (using 90% ethanol and filter sterilized water) Tissue Tearor (Biospec Products, Bartlesville, OK). Most Probable Number (MPN) analyses were performed on individual oyster homogenate and water samples as described in Schuster *et al.* (45). In brief, samples were serially diluted tenfold into Alkaline Peptone Water (APW) and incubated at 37°C for 16 hours, and the tubes scored by turbidity. To positively identify the presence of *V. parahaemolyticus*, 1.0mL of each turbid dilution was pelleted, and the DNA obtained by organic extraction (46). The DNA was subjected to a quantitative q-PCR based MPN as described below, to determine whether each turbid dilution was positive for *V. parahaemolyticus*. The microbiome was recovered from water samples following centrifugation in a 5810R centrifuge (Eppendorf, Hamburg, Germany) at 4,000 rpm and the supernatant discarded. The water bacterioplankton pellet and un-enriched oyster homogenate not immediately used for MPN analysis were frozen at −80°C.

### 4.2 MPN/g enumeration

MPN tubes were scored as positive for *V. parahaemolyticus* by detection of the thermolabile hemolysin gene (*tlh*) with q-PCR (47). The reaction contained 1x iQ Supermix SYBR Green I (Bio-Rad, Hercules, CA) and 2μ L of the DNA template in a final volume of 25μ l. An iCycler with the MyiQ Single Color Real-Time PCR Detection system with included software (Bio-Rad, Hercules, CA, USA) was used with the published cycling parameters (48). A melting curve was performed to ensure positive detection of the correct amplicon compared to a control DNA sample (*V. parahaemolyticus* F11-3A). MPN tubes were scored as positive or negative based on whether q-PCR starting quality values were below (negative) or above (positive) the threshold value determined by the standard curve using purified F11-3A and water blank with iCycler software. The *V. parahaemolyticus* MPN/g was calculated for each oyster according the FDA BAM (24) and grouped by high, medium, or low abundance level based on 10-fold differences in MPN/g.

### 4.3 16s rDNA marker preparation

DNA was isolated from archived oyster homogenates. The homogenates were thawed on ice for 10 minutes, the top ∼ 1cm was aseptically removed and discarded, and 1.0g of each oyster homogenate was aseptically collected. The entire bacterioplankton pellet was used for the water samples. The total bacterial DNA was extracted using the E.Z.N.A. Soil DNA Kit (Omega Bio-Tek, Norcross, GA, USA) following standard protocols for Gram-negative and -positive bacterial isolation.

The V2 to V3 region of 16s rRNA gene (250bp) was amplified from each individual sample in triplicate using PCR with standard 16s F8 (5’ – AGTTTGATCCTGGCTCAG – 3’) with GS FLX Titanium Primer A (5’ – CGTATCGCCTCCCTCGCGCCATCAG – 3’) and R357 (5’ – CTGCTGCCTYCCGTA – 3’) with Primer B (5’ – CTATGCGCCTTGCCAGCCCGCTCAG – 3’), with each pair of corresponding forward and reverse primer sets having a unique 6bp MID tag (48). The PCR reaction containing 45μ L Platinum PCR Supermix (Invitrogen, Carlsbad, CA, USA), 3μ L of sample DNA, and 2μ L molecular grade water, was ran in an iCycler thermocycler (Bio-Rad, Hercules, CA, USA) at the following conditions: 94°C for 90 seconds; 30 cycles of 94°C for 30 seconds, 50.7°C for 45 seconds, 72°C for 30 seconds; and 72°C for 3 minutes. The triplicate samples were combined and then purified using the MinElute PCR Purification Kit (Qiagen, Valencia, CA, USA) following standard protocols. Each purified sample was visualized on a 1.2% agarose gel to ensure purity and quality including expected amplicon size.

A 10ng/mL multiplexed sample was prepared for the Roche Genome Sequencer FLX System using Titanium Chemistry (454 Life Sciences, Branford, CT, USA). The DNA concentration for each sample was quantified using a NanoDrop 2000c (Thermo Scientific, Wilmington, DE, USA) and pooled with equal proportions of the twenty oyster and two water samples. The pooled mixture was purified using the AMPure XP Purification Kit (Beckman Coulter Genomics, Danvers, MA, USA) by manufacturers protocols, with the final samples suspended in 20uL elution buffer EB from the MinElute PCR Purification Kit (Qiagen, Valencia, CA, USA). The pooled tagged single-stranded pyrosequencing library underwent fusion PCR and pyrosequencing using a Roche 454 FLX Pyrosequencer (454 Life Sciences, Branford, CT, USA) according to the manufacturer instructions at the University of Illinois W.M. Keck Center High-Throughout DNA Sequencing Center.

### 4.4 Community analysis

The forward 454 pyrosequencing reads were quality filtered and denoised to reduce erroneous PCR and sequencing errors using FlowClus, setting zero primer and barcode mismatches, a minimum sequence length of 200, zero ambiguous bases and seven homopolymers allowed before truncation, a minimum average quality score of 25, and k=5 for the flow value multiple (49). These sequences were then further filtered and clustered with mothur 1.22.0 (50). The mothur workflow followed the 454 SOP accessed September 2014 (50) with some modifications. The pre-clustering step was performed permitting one difference.Chloroplasts were retained, as cyanobacteria have previously been identified as part of the oyster microbiome (18, 23). The Greengenes 13.8 (51) reference database was used to assign taxonomy to OTUs. After removing singleton OTUs, mothur 1.33.0 (50) was used to generate a distance matrix, pick representative OTUs, and create a phylogenetic tree using clear-cut 1.0.9 (52) for determining alpha diversity.

Rarified alpha diversity measurements were calculated with QIIME 1.8 (53) to determine both the within-sample diversity and sequencing depth using whole-tree phylogenetic diversity (PD) calculated with ten iterations of 100 reads added at each rarefaction step, up to 75% of the sample with the highest number of reads (Supplemental Table 1). The distribution of OTUs between sampling sites and substrates was determined with Venny 1.0 (54). Patterns in abundance in phyla-level classifications in all samples were revealed with a dual-hierarchical clustering performed with JMP 12 (SAS Institute Inc., Cary, North Carolina, USA) for log-transformed percent abundance using both standardized and unstandardized average linkage. Weighted and normalized Fast Unifrac (55), which uses all levels of taxonomic assignment to create a distance matrix and groups samples based on similarity, was used to perform beta diversity clustering and jackknife analyses for samples, jackknifing at 1000 permutations at 75% of the sample with the lowest number of reads. LEfSe (56), PICRUSt (30) and STAMP (57) were all used at default settings, to determine taxonomic and profile similarities between sample groups, and calculate statistical significance, respectively, pre-normalizing samples to 1M in LEfSe.

To compare the sequenced-based abundance of *V. parahaemolyticus* to abundance quantified with the culture-based MPN method, all quality-filtered, de-noised reads were aligned to the region of *V. parahaemolyticus* strain RIMD 2210633 (GCA_000196095.1) that would be amplified by the F8-R357 primer pair at 99.0% with PyNast (58) through QIIME 1.8 (54). The identity of matching sequences was confirmed with BLAST (59). Environmental and nutrient conditions per each site were assessed from the NOAA National Estuarine Research Reserve System (http://nerrs.noaa.gov/) that measures conditions every 15 minutes.

Environmental data used in the statistical analyses were collected as part of this study and the Great Bay National Estuarine Research Reserve (GBNERR) System Wide Monitoring Program (SWMP). Water temperature, salinity, dissolved oxygen, pH, and turbidity were measured and downloaded from YSI datasondes deployed at the study sites from April-December with 15-minute readings. In addition, grab samples collected monthly by GBNERR SWMP were analyzed for chlorophyll *a*, orthophosphate, ammonium, nitrate-nitrite and total dissolved nitrogen (<u>http://cdmo.baruch.sc.edu/get/export.cfm</u>) Short-term environmental conditions, including temperature, salinity, dissolved oxygen, pH, and turbidity were averaged for the 12 hours prior to sampling. Long-term nutrient patterns were assessed by averaging all nutrient analysis data from 2007-2013. The fieldwork performed in this study did not involve endangered or protected species.

## 5. Acknowledgements

We thank C Ellis and A Tyzik for assistance with MPNs, J Jarett and J Gaspar for assistance with bioinformatics software, and W. K. Thomas for discussions. Partial funding for this was provided by the USDA National Institute of Food and Agriculture Hatch NH00574, NH00609 (accession 233555) and NH00625 (accession 1004199). Additional funding provided by the National Oceanic and Atmospheric Administration College Sea Grant program and grants R/CE-137, R/SSS-2, R/HCE-3. Support also provided through the National Institutes of Health 1R03AI081102-01, and National Science Foundation EPSCoR IIA-1330641.This is Scientific Contribution Number 2717 for the New Hampshire Agricultural Experiment Station.

---

**Supplemental Table 1.** Reads available for analysis following quality filtering.

| Oyster | NI | OR |
| --- | --- | --- |
| 1 | 14786 | 34420 |
| 2 | 17900 | 25737 |
| 3 | 17478 | 36390 |
| 4 | 31802 | 25115 |
| 5 | 12500 | 47231 |
| 6 | 10390 | 33229 |
| 7 | 21227 | 24765 |
| 8 | 17670 | 34639 |
| 9 | 21719 | 9672 |
| 10 | 15468 | 22802 |
| Water | 6495 | 397 |

## REFERENCES

(1) Newton AE, Kendall M, Vugia DJ, Henao OL, Mahon BE. Increasing rates of vibriosis in the United States, 1996–2010: Review of surveillance data from 2 systems. Clin Infect Dis. 2012; 54(5): S391–S395.

(2) FDA. Appendix 5: FDA and EPA Safety Levels in Regulations and Guidance. Available: http://www.fda.gov/downloads/Food/GuidanceRegulation/UCM252448.pdf. Accessed 18 December 2015.

(3) Takemura AF, Chien DM, Polz MF. Associations and dynamics of Vibrioaceae in the environment, from the genus to the population level. Front Microbiol. 2014; 5: 38.

(4) Larsen AM, Rikard FS, Walton WC, Arias CR. Temperature effect on high salinity depuration of Vibrio vulnificus and V. parahaemolyticus from the easter oyster (Crassostrea virginica). 2015. Int J Food Microiol 192:66–71.

(5) Parveen S, Jahncke M, Elmahdi S, Crocker H, Bowers J, White C, Gray S, Morris AC, Brohawn K. High salinity relaying to reduce Vibrio parahaemolyticus and Vibrio vulnificus in Chesapeake Bay oysters (Crassostrea virginica). 2017. J Food Sci. 82:484–491.

(6) Yu JW. Incidence, abundance, post-harvest processing and population diversity of pathogeni Vibrios in oysters from the Great Bay Estuary. University of New Hampshire, ProQuest Dissertations Publishing, 2011. 1507840

(7) Main CR, Salvitti LR, Whereat EB, Coyne KJ. Community-level and species-specific associations between phytoplankton and particle-associated *Vibrio* species in Delaware’s inland bays. Appl Environ Microbiol. 2015; 81(17): 5703–5713

(8) Urquhart EA, Jones SH, Yu JW, Schuster BM, Marcinkiewicz AL, Whistler CA, et al. Environmental conditions associated with elevated risk conditions for *Vibrio parahaemolyticus* in the Great Bay Estuary, NH. PLoS ONE. 2015; 11(5): e0155018.

(9) Matz C, Nouri B, McCarter L, Martinez-Urtaza J. Acquired type III secretion system dertermines environmental fitness of epidemic Vibrio parahaemolyticus in the interaction with bacteriouvorous protists. 2011. PLoS One 6:e20275

(10) Lokmer A, Goedknegt MA, Thieltges DW, Fiorentino D, Kuenzel S, Baines JF, Wegner KM. Spatial and temporal dynamics of Pacific oyster hemolymph microbiotia across multiple scales. 2016 7:13678

(11) Givens CE, Bowers JC, DePaola A, Hollibaugh JT, Jones JL. Occurrence and distribution of *Vibrio vulnificus* and *Vibrio parahaemolyticus* – potential roles for fish, oyster, sediment, and water. Letters in Appl Microbiol. 2014; 58(6): 503–510.

(12) Olafsen JA, Mikkelsen HV, Giaever HM, Hansen GH. Indigenous bacteria in hemolymph and tissues of marine bivalves at low temperatures. Appl Environ Microbiol 1993; 59(6): 1848-1854.

(13) Pujalte MJ, Ortigosa M, Maciá MC, Garay E. Aerobic and facultative anaerobic heterotrophic bacteria associated to Mediterranean oysters and seawater. Int Microbio. 1999; l2(4): 259–266.

(14) Volety AK, McCarthy SA, Tall BD, Curtis SK, Fisher WS, Genthner FJ. Responses of oyster *Crassostrea virginicia* hemocytes to environmental and clinical isolates of *Vibrio parahaemolyticus*. Aquat Microb Ecol. 2001; 25: 11–20.

(15) Long, RA, Azam F. Antagonistic interactions among marine pelagic bacteria. Appl Enviro Microbiol. 2001; 67(11): 4975–4983.

(16) Rypien KL, Ward JR, Azam F. Antagonistic interactions among coral-associated bacteria. Environ Microbiol. 2009; 12(1): 28–39.

(17) Frydenborg, BR, Krediet CJ, Teplitski M, Ritchie KB. Temperature-dependent inhibition of opportunistic Vibrio pathogens by native coral commensal bacteria. Microb Ecol. 2014; 67(2): 392–401.

(18) Chauhan A, Wafula D, Lewis DE, Pathak A. Metagenomic assessment of the eastern oyster-associated microbiota. Genome Announc. 2014; 2(5): e01083–14.

(19) Chen H, Liu Z, Wang M, Chen S, Chen T. Characterization of the spoilage bacterial microbiota in oyster gills during storage at different temperatures. J Sci Food Agric. 2013; 93(15): 3748–3754.

(20) King GM, Judd C, Kuske CR, Smith C. Analysis of stomach and gut microbiomes of the eastern oyster (*Crassostrea virginica*) from coastal Louisiana, USA. PLoS One. 2012; 7(12): e51475.

(21) Trabal-Fernandez N, Mazon-Suastegui JM, Vazquez-Juarez R, Ascencio-Valle F, Romero J. Changes in the composition and diversity of the bacterial microbiota associated with oysters (*Crassostrea corteziensis, Crassostrea gigas and Crassostrea sikamea*) during commercial production. FEMS Microbiol Ecol. 2013; 88: 69–83.

(22) Wegner KM, Volkenborn N, Peter H, Eiler A. Disturbance induced decupling between host genetics and composition of the associated microbiome. BMC Microbiol. 2013; 13: 252.

(23) Zurel D, Benayahu Y, Or A, Kovacs A, Gophna U. Composition and dynamics of the gill microbiota of an invasive Indo-Pacific oyster in the eastern Mediterranean Sea. Environ Microbiol. 2011; 13(6): 1467–1476.

(24) FDA. Bacteriological analytical manual. Chapter 9. Vibrio. Kaysner, CA and A. DePaola. http://www.fda.gov/Food/FoodScienceResearch/LaboratoryMethods/ucm2006949.htm. Accessed 8 March 2016.

(25) Thomas IV JC, Wafula D, Chauhan A, Green SJ, Gragg R, Jagoe C. A survey of deepwater horizon (DWH) oil-degrading bacteria from the Eastern oyster biome and its surrounding environment. Front Microbiol. 2009; 5: 149.

(26) Avila-Poveda OH, Torres-Ariñ o A, Girón-Cruz DA, Cuevas-Aquirre A. Evidence for accumulation of *Synechococcus elongatus* (Cyanobacteria: Cyanophyceae) in the tissues of the oyster *Crassostrea gigas* (Mollusca: Bivalvia). Tissue Cell. 2014; 46(5): 379–387.

(27) Berg KA, Lyra C, Sivonen K, Paulin L, Suomalainen S, Tuomi P, et al. High diversity of cultivable heterotrophic bacteria in association with cyanobacterial water blooms. ISME J. 2009; 3(3): 314–325.

(28) Levy P-Y, Fenollar F, Stein A, Borrione F, Raoult D. *Finegoldia magna*: a forgotten pathogen in prosthetic joint infection rediscovered by molecular biology. Clin Infect Dis. 2009; 49(8): 1244–1247.

(29) Jordan DC. NOTES: Transfer of *Rhizobioum japonicum* Buchanan 1980 to *Brazyrhizobium* gen. nov., a genus of slow-growing, root nodule bacteria from leguminous plants. Int J Syst Evol Microbiol. 1982; 32: 136–139.

(30) Langille MGI, Zaneveld J, Caporaso JG, McDonald D, Knights D, Reyes JA, et al. Predictive functional profiling of microbial communities using 16S rRNA marker gene sequences. Nat Biotechnol. 2013; 31: 814–821.

(31) Drury B, Rosi-Marshall E, Kelly JJ. Wastewater treatment effluent reduces the adundance and diversity of benthic bacterial communities in urban and suburban rivers. Appl Environ Microbiol. 2013; 79(6): 1897–1905.

(32) Li D, Sharp JO, Drewes JE. Influence of wastewater discharge on the metabolic potential of the microbial community in river sediments. Microb Ecol. 2016; 71(1): 78–86.

(33) Kaufman GE, Bej AK, DePaola A. Oyster-to-oyster variability in levels of *Vibrio parahaemolyticus*. J Food Prot. 2003; 66(1): 125–129.

(34) Klein SL, Lovell CR. The Hot oyster: levels of virulent Vibrio parahaemolyticus strains in individual oysters. 2017 93(2) pii:fiw232

(35) Jones SH, Summer-Brason BW. 1998. Incidence and detection of pathogenic Vibrio sp. in a northern New England estuary, USA. J. Shellfish Res. 17:1665–1669.

(36) Givens CE, Ransom B, Bano N, Hollibaugh JT. Comparison of the gut microbiomes of 12 bony fish and 3 shark species. Mar Ecol Prog Ser. 2015; 518: 209–223.

(37) Ward JE, Shumway SE. Separating the grain from the chaff: particle selection in suspension-and deposit-feeding bivalves. J Exp Mar Biol Ecol. 2004; 300: 83–130.

(38) Birkbeck TH, McHenery JG. Degradation of bacteria by *Mytilus edulis*. Marine Biol. 1982; 72: 7–15.

(39) Murphree RL, Tamplin ML. Uptake and retention of *Vibrio cholerae* O1 in the Eastern oyster, *Crassostrea virginica*. Appl Environ Microbiol. 1995; 61(10): 3656–3660.

(40) Paranjpye RN, Johnson AB, Baxter AE, Strom MS. Role of type IV pilins in persistence of *Vibrio vulnificus* in *Crassostrea virginica* oysters. Appl Environ Microbiol. 2007; 73(15): 5041– 5044.

(41) Srivastava M, Tucker MS, Gulig PA, Wright AC. Phase variation, capsular polysaccharide, pilus, and flagella contribute to uptake of *Vibrio vulnificus* by the Eastern oyster (*Crassostrea virginica*). Environ Microbiol. 2009; 11(8): 1934–1944.

(42) Soizick LG, Robert A, Haifa M, Jacques LP. Shellfish contamination by norovirus: strain selection based on ligand expression?. Clin Virol. 2013; 41(1): 3–18.

(43) Froelich BA, Ringwood A, Sokolova I, Oliver JD. Uptake and depuration of the C- and E-gneotypes of *Vibrio vulnificus* by the Eastern oyster (*Crassostrea virginicia*). Environ Microbiol Rep. 2009; 2(1): 112–115.

(44) Genther FJ, Volety AK, Oliver LM, Fisher WS. Factors influencing *in vitro* killing of bacteria by hemocytes of the Eastern oyster (*Crassostrea virginica*). Appl Enviro Microbiol. 1999; 65(7): 3015–3020.

(45) Schuster BM, Tyzik AL, Donner RA, Striplin MJ, Almagro-Moreno S, Jones SH, et al. Ecology and genetic structure of a northern temperate *Vibrio cholerae* population related to toxigenic isolates. Appl Enviro Microbiol. 2011; 77(21): 7568–7575.

(46) Ausubel F, Brent R, Kingston RE, Moore DD, Seidman JG, Smith JA, et al. Current protocols in molecular biology. New York, N.Y.: Wiley and Sons, Inc.; 1990.

(47) Nordstrom JL, Vickery MCL, Blackstone GM, Murray SL, DePaola A. Development of a multiplex real-time PCR assay with an internal amplification control for the detection of total and pathogenic *Vibrio parahaemolyticus* bacteria in oysters. Appl Enviro Microbiol. 2007; 73(18): 5840–5847.

(48) Liu Z, Lozupone C, Hamady M, Bushman FD, Knight R. Short pyrosequencing reads suffice for accurate microbial community analysis. Nucleic Acids R. 2007; 35(18): e120-.

(49) Gaspar JM, Thomas WK. FlowClus: Efficiently filtering and denoising pyrosequenced amplicons. BMC Bioinformatics 2015; 16: 105.

(50) Schloss PD, Westcott SL, Ryabin T, Hall JR, Hartmann M, Hollister EB, et al. Introducing mothur: open-source, platform-independent, community-supported software for describing and comparing microbial communities. Appl Environ Microbiol. 2009; 75(23): 7537–7541.

(51) DeSantis TZ, Hugenholtz P, Larsen N, Rojas M, Brodie EL, Keller K, et al. Greengenes, a chimera-checked 16S rRNA gene database and workbench compatible with ARB. Appl Environ Microbiol. 2006; 72(7): 5069–5072.

(52) Sheneman L, Evans J, Foster JA. Clearcut: A fast implementation of relaxed neighbor joining. Bioinformatics. 2006; 22(22): 2823–2824.

(53) Caporaso JG, Kuczynski J, Stombaugh J, Bittinger K, Bushman FD, Costello EK, et al. QIIME allows analysis of high-throughput community sequencing data. Nat Meth. 2010; 7(5): 335–336.

(54) Oliveros JC. Venny, an interactive tool for comparing lists with Venn’s diagrams. 2007-2015. http://bioinfogp.cnb.csic.es/tools/venny/index.html.

(55) Lozupone C, Lladser ME, Knights D, Stombaugh J, Knight R. UniFrac: an effective distance metric for microbial community comparison. ISME J. 2011; 5(2): 169–172.

(56) Segata, N, Izard J, Waldron L, Gevers D, Miropolsky L, Garrett WS, et al. Metagenomic biomarker discovery and explanation. Genome Biol. 2011; 12(6): R60.

(57) Parks DH, Tyson GW, Hugenholtz P, Beiko RG. STAMP: statistical analysis of taxonomic and functional profiles. Bioinformatics. 2014; 30(1): 3123–3124.

(58) Caporaso JG, Bittinger K, Bushman FD, DeSantis TZ, Andersen GL, Knight R. PyNAST: A flexible tool for aligning sequences to a template alignment. Bioinformatics. 2010; 26(2): 266– 267.

(59) Altschul SF, Gish W, Miller W, Myers EW, Lipman DJ. Basic local alignment search tool. J Mol Biol. 1990; 215:403–410.

